# A redox switch allows binding of Fe(II) and Fe(III) ions in the cyanobacterial iron binding protein FutA from *Prochlorococcus*

**DOI:** 10.1101/2023.05.23.541926

**Authors:** Rachel Bolton, Moritz M. Machelett, Jack Stubbs, Danny Axford, Nicolas Caramello, Lucrezia Catapano, Martin Malý, Matthew J. Rodrigues, Charlotte Cordery, Graham J. Tizzard, Fraser MacMillan, Sylvain Engilberge, David von Stetten, Takehiko Tosha, Hiroshi Sugimoto, Jonathan A.R. Worrall, Jeremy S. Webb, Mike Zubkov, Simon Coles, Eric Mathieu, Roberto A. Steiner, Garib Murshudov, Tobias E. Schrader, Allen M. Orville, Antoine Royant, Gwyndaf Evans, Michael A. Hough, Robin L. Owen, Ivo Tews

## Abstract

The marine cyanobacterium *Prochlorococcus* is a main contributor to global photosynthesis, whilst being limited by iron availability. Cyanobacterial genomes typically encode two different types of FutA iron binding proteins: periplasmic FutA2 ABC transporter subunits bind Fe(III), while cytosolic FutA1 binds Fe(II). Owing to their small size and their economized genome *Prochlorococcus* ecotypes typically possess a single *futA* gene. How the encoded FutA protein might bind different Fe oxidation states was previously unknown. Here we use structural biology techniques at room temperature to probe the dynamic behavior of FutA. Neutron diffraction confirmed four negatively charged tyrosinates, that together with a neutral water molecule coordinate iron in trigonal bipyramidal geometry. Positioning of the positively charged Arg103 side chain in the second coordination shell yields an overall charge-neutral Fe(III) binding state in structures determined by neutron diffraction and serial femtosecond crystallography. Conventional rotation X-ray crystallography using a home source revealed X-ray induced photoreduction of the iron center with observation of the Fe(II) binding state; here, an additional positioning of the Arg203 side chain in the second coordination shell maintained an overall charge neutral Fe(II) binding site. Dose series using serial synchrotron crystallography and an XFEL X-ray pump-probe approach capture the transition between Fe(III) and Fe(II) states, revealing how Arg203 operates as a switch to accommodate the different iron oxidation states. This switching ability of the *Prochlorococcus* FutA protein may reflect ecological adaptation by genome streamlining and loss of specialized FutA proteins.

**Significance Statement:** Oceanic primary production by marine cyanobacteria is a main contributor to carbon and nitrogen fixation. *Prochlorococcus* is the most abundant photosynthetic organism on Earth, with an annual carbon fixation comparable to the net global primary production from agriculture. Its remarkable ecological success is based on the ability to thrive in low nutrient waters. To manage iron limitation, *Prochlorococcus* possesses the FutA protein for iron uptake and homeostasis. We reveal a molecular switch in the FutA protein that allows it to accommodate binding of iron in either the Fe(III) or Fe(II) state using structural biology techniques at room temperature and provide a plausible mechanism for iron binding promiscuity.

## Introduction

Iron is the fourth most abundant element in the Earth’s crust (1). However, because of the poor solubility, primary production in large oceanic and freshwater environments is limited by iron uptake (2). In oxygenated aqueous environments, iron predominantly exists in Fe(III) oxyhydroxides (3) with a solubility of 10^-18^ M (4) and consequently precipitates to severely limit bioavailability (5). Marine phytoplankton require iron in the photosynthetic electron transport chain (6) and in the nitrogenase enzyme (7, 8); thus, iron availability directly limits photosynthesis (9) and nitrogen fixation (10).

Cyanobacteria of the *Prochlorococcus* genus are able to fix four gigatons of carbon per annum, which is comparable to the net primary production of global agriculture (11). *Prochlorococcus* bacteria dominate bacterial populations in tropical and subtropical oligotrophic ocean regions (12). One of the factors for ecological success is the exceptional ability of this bacterium to thrive in low nutrient waters (13). Adaptation includes reduction in size to 0.5 – 0.7 μm, making *Prochlorococcus* not only the most abundant but also the smallest photosynthetic organism on Earth (14). Reduction in size maximizes the surface-area-to-volume ratio for metabolic efficiency, to a tradeoff of genome reduction, and *Prochlorococcus* maintains the smallest genome (1.6-2.7 Mb) known for any free-living phototroph (15).

Typically, cyanobacteria harbor multiple iron uptake systems (16). In the common TonB transport system, organic ligands (siderophores) are used to solubilize iron (17). The majority of the *Prochlorococcus* species lack genes for siderophore biosynthesis (18, 19); instead, the bacterium relies on the Fut ABC transporter for iron uptake (20). Here, specialized periplasmic proteins sequester elemental iron (16); FutA2 is such a substrate binding protein (SBP) that binds Fe(III) to deliver it to the Fut ABC transporter (21, 22). A functional homologue of FutA2 is the cytosolic protein FutA1 that binds Fe(II) and protects the photosystem against oxidative stress (23-25); however, FutA1 has also been shown to bind Fe(III) (21, 26). We have previously reported dual localization and function for the single FutA protein of the marine cyanobacterium *Trichodesmium* (27), suggesting it can bind both iron species. Similarly, *Prochlorococcus* harbors a single *futA* gene (20), we wanted to understand whether and how a single FutA protein can bind both iron species, and how redox plasticity was structurally encoded.

It is challenging to obtain crystallographic models without alteration of the metal sites, since site-specific damage occurs extremely quickly and at very low doses (28), particularly for iron (29, 30). Indeed, the FutA structure determined from a conventional diffraction experiment on an X-ray home source reported here represented the photo-reduced, Fe(II) binding state, corroborated by spectroscopic evidence. A serial femtosecond crystallography approach (SFX) using an XFEL source and a complementary neutron diffraction approach were required to avoid the manifestations of X-ray induced photoreduction in order to determine the Fe(III) state and give protonation states of iron coordinating amino acid side chains. Using a fixed-target silicon chip system for crystal delivery (31) at both synchrotron and XFEL radiation sources, we studied the transition between Fe(III) to Fe(II) states whilst making use of the effects of X-ray induced photoreduction, varying dose and time. The resulting protein structures support a dual binding mode for iron and give insight into protein adaptation to evolutionary pressures.

## Results

### The structure of FutA

The crystallographic X-ray structure of FutA was determined from a single crystal to 1.7 Å resolution, using a standard rotation protocol with the crystal in a sealed capillary at a home source setup (**Table S1**). Substrate binding domains such as FutA can be classified based on overall fold and *Prochlorococcus* FutA classifies as "D type” substrate binding protein. The N-terminal (amino acids 1-98 and 232-280, light grey) and C-terminal domains (amino acids 99-231 and 281-314, dark grey) are highlighted in **Fig. 1A**.

**Figure 1.**
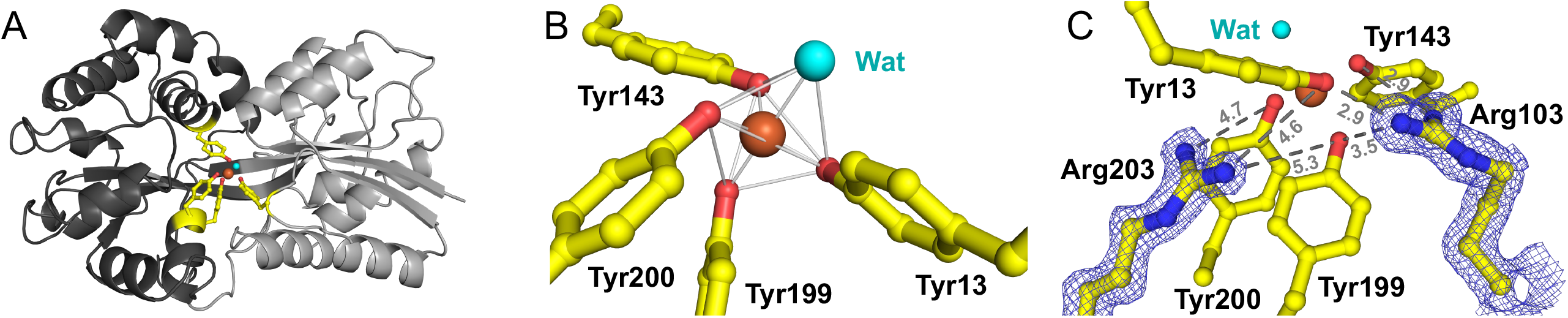
The Fe(II) state FutA structure from an X-ray home source determined to 1.7 Å resolution. (*A*) FutA has a bi-lobal structure with the substrate binding cleft between the N-terminal (light grey) and C-terminal domains (dark grey). Amino acid side chains contributing to iron binding are shown in stick representation (yellow). (*B*) Trigonal bipyramidal coordination of the iron, with Tyr199 and a solvent molecule as axial ligands. (*C*) The two arginine side chains of Arg103 and Arg203 are in a second coordination shell, shown here with refined density (2Fobs – Fcalc, blue map, contoured at 1.5 σ). Color coding is yellow for carbon, red for oxygen, blue for nitrogen, orange for iron, with the solvent molecule in light blue.

The substrate-binding cleft bears the iron-binding site that is open to the surrounding solvent. The four tyrosine side chains of Tyr13 from N-terminal and Tyr143, Tyr199 and Tyr200 from C-terminal domains coordinate the iron, **Fig. 1B**, in this Class IV substrate binding protein (32). The trigonal bipyramidal coordination involves Tyr13, Tyr143 and Tyr200 to form the trigonal plane with iron at its center, while Tyr199 and a coordinating solvent molecule are the axial ligands.

Interestingly, the structure reveals a positioning of two arginine side chains, Arg103 and Arg203, in a second shell around the iron binding site, **Fig. 1C**. One might assume the tyrosine side chains are negatively charged tyrosinates, and arginine side chains would each provide a positive charge, with a neutral solvent molecule. To understand the charge state, we used spectroscopy and confirmed protonation states using neutron diffraction.

### Determination of the Fe(III) iron binding state by spectroscopy

A refolding protocol in presence of iron sulfate was used to purify FutA. The burgundy red color of the purified protein that can readily be bleached by excess sodium dithionite likely resulted from the ligand to metal charge transfer (LMCT) bands between the tyrosinate residues coordinating the Fe(III) ion, **Fig. 2A**.

**Figure 2.**
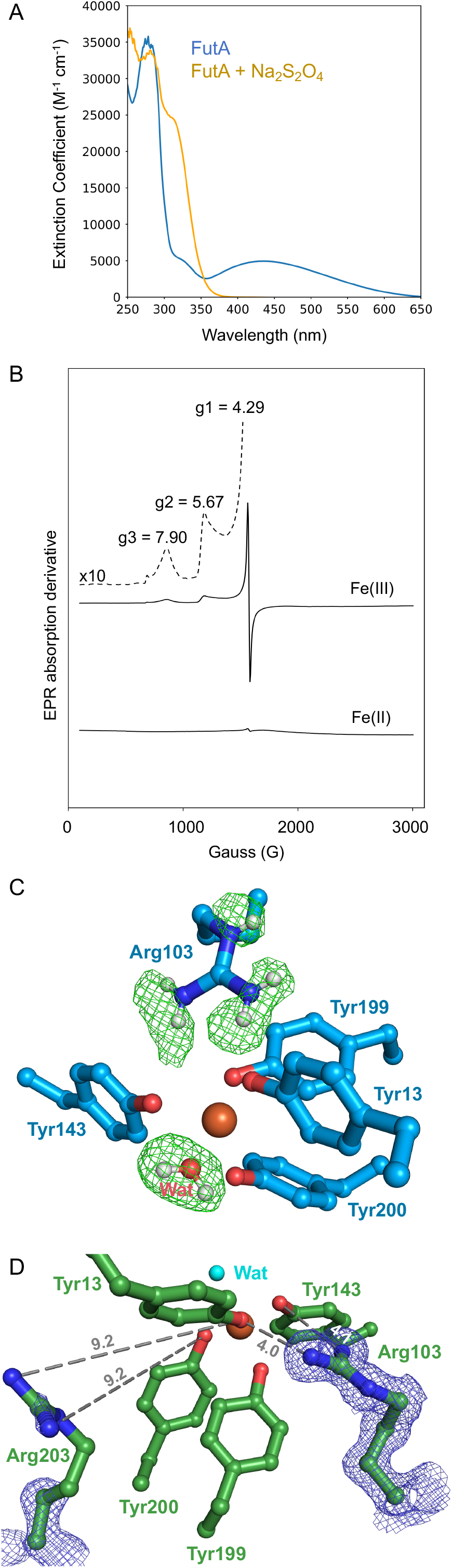
The FutA Fe(III) state characterized by UV-vis and EPR spectroscopy, neutron diffraction and serial femtosecond crystallography. (*A*) The UV-vis spectrum of recombinantly produced and purified FutA (blue) shows an absorbance maximum at 438 nm, consistent with Fe(III) bound to FutA. The peak at 438 nm disappears after addition of 10-fold molar excess sodium dithionite; the absorbance maximum at 315 nm indicates free sodium dithionite (yellow). (*B*) EPR spectrum of purified and sodium dithionite reduced FutA. The peaks observed were: g1 = 4.29 g, g2 = 5.67 g, g3 = 7.9 g. (*C*) The positive nuclear density in the neutron diffraction crystal structure (green mesh, Fobs – Fcalc omit map at 3σ, 2.1 Å resolution) indicates sites that have undergone hydrogen-deuterium exchange, showing an oriented water as axial ligand (refined deuterium fraction > 0.80). Arg103 fully protonated and positively charged, while the four tyrosine side chains do not show difference density, suggesting they are negatively charged tyrosinates. (*D*) The SFX crystal structure shows that the side chain of Arg203 is not oriented towards the binding site and does not engage in polar interactions (similar to the neutron diffraction structure **Fig. S1**). Carbons shown blue (neutron diffraction) or green (SFX), heteroatoms colored as in **Fig. 1**.

The electron paramagnetic resonance (EPR) spectrum of purified FutA shows a sharp signal at a g-value of 4.29, **Fig. 2B**. This signal is indicative of a |±3/2> doublet from a 3d^5^, high-spin (S = 5/2) isotropic system (E/D ≈ 1/3), consistent with an Fe(III) ion bound to FutA (33). The weaker signals (g = 5.67, g = 7.90) derive from either |±1/2> ground state transitions or from |±3/2> resonances from rhombic species of the Fe(III) iron. However, given the very high transition probabilities for the g = 4.29 signal compared to the lower transition probability for ground state or anisotropic species, the latter resonances likely represent a significant fraction of the total spins in the sample. Excess of sodium dithionite leads to the loss of the EPR signal, **Fig. 2B**. This could result from loss of iron binding and reduction in solution, or reduction of Fe(III) iron to a colorless and 3d^6^ EPR-silent (probably S=2) Fe(II) state within the active site.

### Protonation state of Fe(III) coordinating residues as determined by neutron diffraction

We determined the crystallographic structure of FutA by neutron diffraction to 2.1 Å resolution (**Tables S1 & S2**). Positive density in the neutron Fo-Fc omit map indicates sites of successful hydrogen-deuterium exchange. The lack of deuterium on the iron coordinating Tyr13, Tyr143, Tyr199 and Tyr200 suggests these residues are tyrosinates, **Fig. 2C**. The nuclear density for the metal-bound solvent is consistent with neutral water. Arg203 is not engaged in any interactions and does not contribute to the second shell (**Fig. S1**), in contrast to the X-ray structure, **Fig. 1**. However, side chain of Arg103 in the second shell is fully protonated and positively charged, thus together with the four negatively charged tyrosinates Fe(III) binding results in an overall charge balanced binding site.

### The Fe(III) iron state structure determined by serial femtosecond crystallography (SFX)

The SFX experiment used short (10 fs), high-intensity X-ray pulses from the SACLA XFEL to provide diffraction patterns that are collected before the crystal is destroyed (34). It has been shown that data can be recorded free of the effects of radiation induced changes as long as sufficiently short pulses (<20 fs) are used (35). Crystallization conditions were optimized to obtain microcrystal slurries suitable for SFX, as described by us previously (36). For data collection, crystals of approximately 20 x 7 x 7 μm^3^ were applied onto a fixed-target silicon chip, with the final dataset merged from three chips (**Table S1**).

SFX and neutron diffraction structures are similar (see comparison in SI), with the Arg103 side chain contributing to the second shell, but the side chain of Arg203 pointing away from the binding site, **Fig. 2D**. EPR data, neutron diffraction and SFX agree and are consistent with iron binding in the Fe(III) state. In turn, this suggests that the structure determined from the X-ray home source with the Arg203 side chain pointing towards the binding site as shown in **Fig. 1** may represent the Fe(II) state.

### Characterization of X-ray induced photoreduction of Fe(III) FutA

The home source rotation experiment might either fortuitously have captured the reduced state, or this observation had resulted from X-ray induced photoreduction of Fe(III) to Fe(II). Photoreduction was highly likely, considering the bleaching of the burgundy-red appearance in the X-ray exposed area of the crystal during data collection. We thus went on to characterize the effect of X-ray exposure using *in crystallo* optical spectroscopy (37).

The electronic absorption peak (λmax = 438 nm) corresponding to the Fe(III) iron (38) progressively decays on incident X-ray irradiation at a synchrotron beamline, **Fig. 3A**. As X-rays induce light-absorbing chemical species in the solvent that overlap with the Fe(III) iron specific signal, the 620 nm wavelength was chosen to minimize the effect of this artefact and characterize photoreduction of the iron center, plotting absorbance against accumulated radiation dose, **Fig. 3B**. Measuring five different crystals, we determined a half-photoreduction dose of 128 ± 21 kGy; the dose at which 80% of the molecules had been photoreduced was 204 ± 27 kGy.

**Figure 3.**
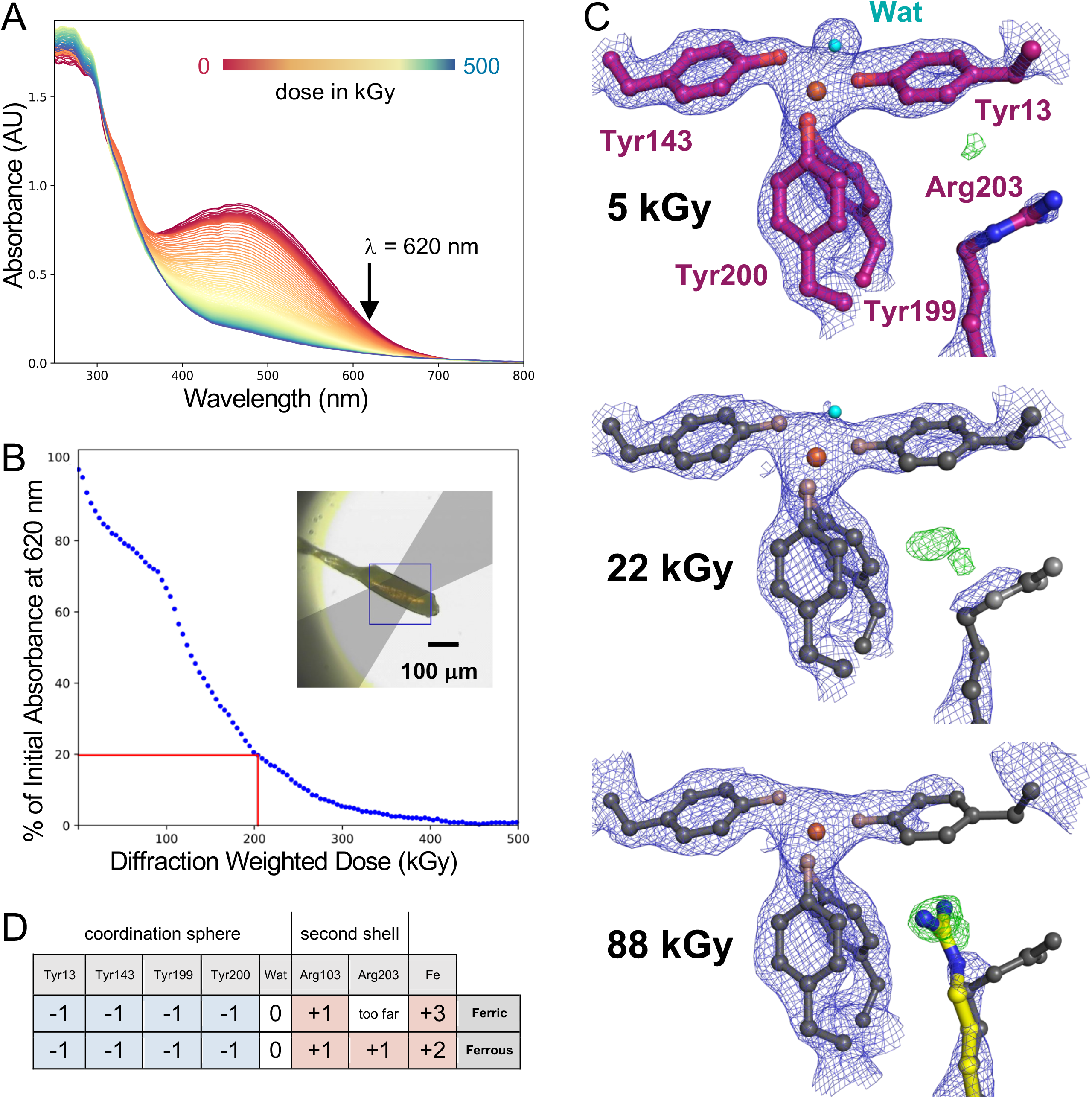
X-ray induced photoreduction of FutA characterized by spectroscopy and SSX. (*A*) Successive UV-vis absorption spectra collected *in crystallo* plotted for a FutA crystal during X-ray exposure, from 0 kGy (red) to 500 kGy (blue). Photoreduction was monitored at a wavelength of 620 nm (arrow). (*B*) Evolution of the normalized absorbance at 620 nm, collected on a single crystal. In the example shown, 80% of the signal was lost at 204 ± 27 kGy (red lines). Inset: geometry of the experiment. The light path for the spectroscopic measurement is indicated in grey. (*C*) SSX dose series at RT. Top: refined structure at 5 kGy (carbon atoms shown in purple; 2Fo-Fc density in blue contoured at 1.5 σ, Fo-Fc in green contoured at 3 σ). Pronounced difference density is seen at 22 kGy and 88 kGy, suggesting Arg203 takes an alternative conformation, as indicated by overlay with the conformation seen in the Fe(II) state determined from the home source (Arg203 carbons shown in yellow for the 88 kGy dose point). Heteroatoms colored as in **Fig. 1**. (*D*) Charges of amino acids contributing to the coordination sphere and second shell for the Fe(III) and Fe(II) binding states, assuming an overall neutral state of the binding site.

### Tracking of X-ray induced photoreduction from an SSX dose series

A fixed target serial synchrotron crystallography (SSX) approach described by us previously (31) was used, as it is well suited for low dose investigations. A series of ten images was taken from each microcrystal, where each image incrementally increases the dose, allowing us to follow structural changes of the FutA iron complex in response to X-ray induced photoreduction.

Two different dose series with dose increments of 5 kGy and 22 kGy are reported (**Tables S3 & S4**). Images corresponding to each dose point are merged to provide a series of datasets corresponding to these dose points. The isomorphous difference density indicates an alternative conformation for Arg203. The feature is readily visible at 22 kGy and strongest at 88 kGy, **Fig. 3C**. Indeed, overlay with the conformation observed in the home source structure, **Fig. 1C**, shows that both structures are similar, suggesting the photoreduced state was observed in either case.

### An XFEL X-ray pump-probe (XRPP) approach captures the transition between Fe(III) and Fe(II) states

We designed a novel serial femtosecond crystallography experiment where a first pulse, attenuated using a sapphire wafer mounted on a fast flipper, was followed by a second, unattenuated pulse (**Fig. S2)**. Using SACLA’s repetition rate of 30 Hz, the 10 fs pump and probe were spaced 33 ms apart. While several different levels of attenuation were explored, data for a 350 kGy pump (94% attenuated) yielded structural changes consistent with photoreduction. Interestingly, in contrast to the SSX series, **Fig. 3C**, this experiment preserved the iron coordinating water that was clearly resolved in electron density, **Fig. 4**, consistent with penta-coordinated Fe(II) iron. Ensuing refinement confirms presence of the alternative conformation of Arg203 (**Fig. S3**). For the high occupancy state of Arg203 with the guanidino group closest to the iron center, distances were 4.8 Å between the η1 amide of Arg203 and the phenolate oxygen of Tyr200, and 4.6 Å between the η2 amide of Arg203 and the alkoxy group of Tyr13. The XRPP experiment thus induced specific alteration(s) and created the FutA Fe(II) state *in situ*.

**Figure 4.**
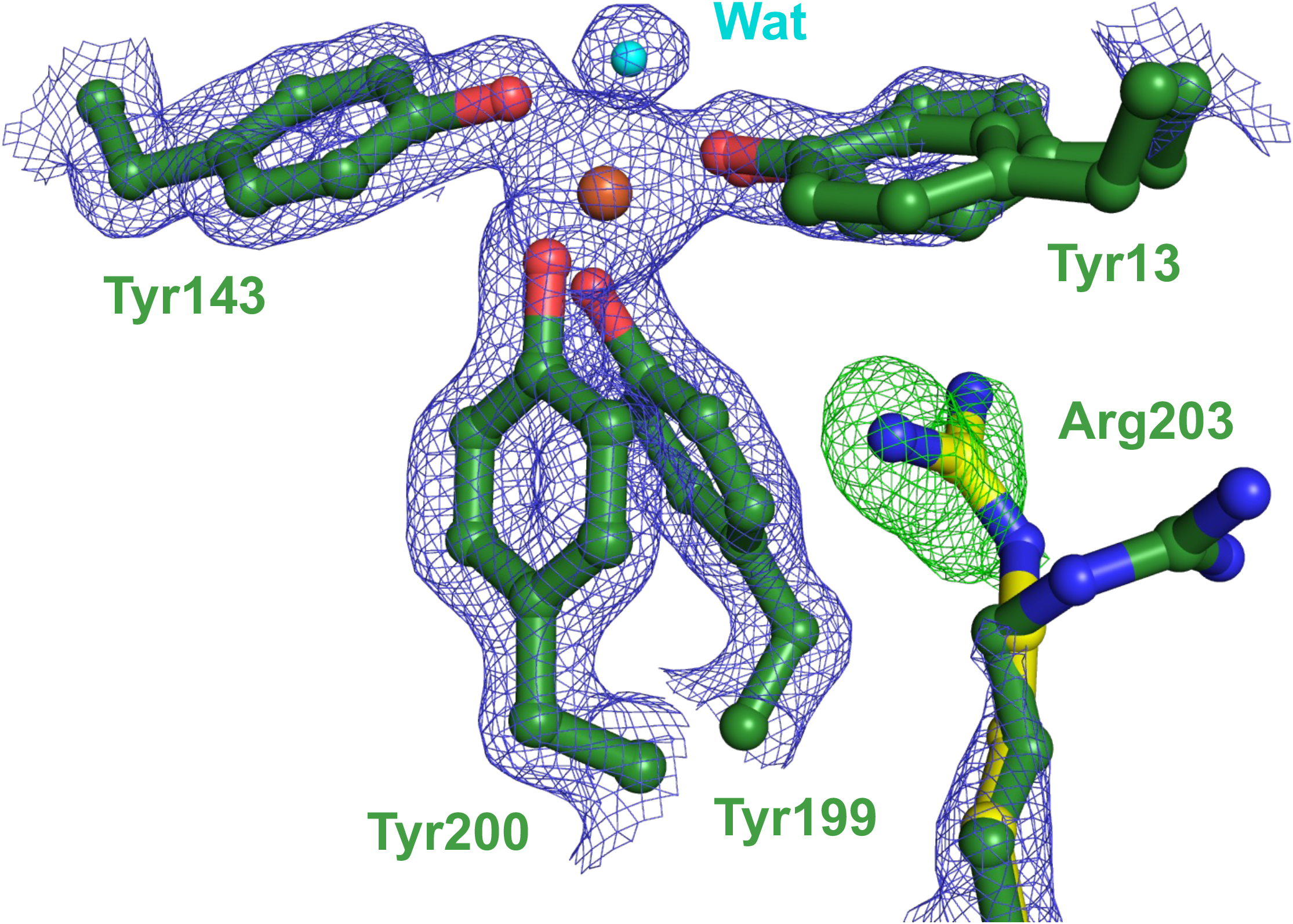
SFX X-ray pump probe experiment. The model of the Fe(III) iron state determined SFX (compare **Fig. 2D**) was used in refinement against an SFX probe dataset, collected after a 350 kGy pump. Refined electron density shows Tyr13 in a double conformation, but limited density for the Arg203 guanidino group (2Fo-Fc, blue, 1.5 σ); however, difference density (Fo-Fc, green, 3 σ) suggests that Arg203 takes an alternative conformation similar to the conformation observed in the Fe(II) state determined from the home source (Arg203 carbons shown in yellow). Heteroatoms colored as in **Fig. 1**.

## Discussion

The adaptation of the marine cyanobacterium *Prochlorococcus* is a remarkable story of ecological success, making this photosynthetic organism the most abundant on earth. Two factors are particularly important, the ability to survive under limiting nutrient conditions and physical size reduction where both factors put evolutionary pressure on the iron uptake system of the bacterium (13, 14). This study addresses the challenge of how a single gene product, FutA, can bind both Fe(III) and Fe(II) iron.

The structural analyses reported at ambient (room) temperature allow delineating a plausible mechanism for iron binding in two different oxidation states, showing how FutA Arg203 operates as a switch between states. The side chain of this residue is not engaged in polar contacts in the Fe(III) states, which is hinting at its intrinsic dynamics, allowing it to be recruited and engage in interaction with the iron center and contribute a balancing charge in the Fe(II) state, **Fig. 3D**.

X-ray crystallographic study of RedOx active metallo-proteins is challenging as X-ray induced photoreduction can occur. Transition metals are particularly sensitive to specific radiation damage (28, 39), and observation of the FutA Fe(III) state required SFX / neutron diffraction. Changes in the oxidation state induced by X-rays were previously documented for doses as low as 30 - 20 kGy (30, 40, 41). For Fe(III), we show that the half-point for photoreduction in FutA corresponds to a dose of 128 +/- 21 kGy, as shown by spectroscopic analysis, **Fig. 2B**.

We exploited the effects of X-ray induced photoreduction to study the transition between Fe(III) and Fe(II) states, using a SSX dose series and an SFX X-ray pump probe setup, both allowing us to map conformational changes at ambient temperatures (42). The major difference observed between these approaches occurred for density of the coordinating water, which disappeared with accumulating dose in SSX, **Fig. 3C**, while the SFX XRPP approach preserved the electron density, **Fig. 4**. The reported doses delivered were 350 kGy in a 10-femtosecond pulse in the SFX XRPP pump which was 3.2 and 4 times higher than the doses used in the home source and SSX experiments, respectively, both of which used continuous X-ray exposure which could lead to difference in heat load. Beam sizes were also different, with 300 micron for home source, 10 micron for SSX and 1.5 micron for SFX experiments, providing an important difference for photo-electron escape (43). Further work is needed to understand radiation chemistry arising from these different conditions (see additional discussion in SI).

Discovery of a mechanism to bind two different iron oxidation states prompted us to revisit homologues of the FutA iron binding protein, and we found that a similar switch may exist for the iron binding protein FbpA from *Thermus thermophilus* with structures in two states reported (**Fig. S4**). *Synechocystis* has two specialized iron binding proteins, with FutA2 being assigned a Fe(III) binding function in the oxidative environment of the periplasm, while FutA1 binds Fe(II) iron favored under reducing conditions in the cytosol. For these proteins, conservation of the arginine residue equivalent *Prochlorococcus* Arg203 (**Fig. S5**) may relate to biological ability to bind iron at different oxidation states, as discussed in supplementary text.

## Conclusion

Structures with iron bound in different oxidation states help explain how the intrinsic structural plasticity of FutA accommodate Fe(II) as well as Fe(III) iron species. Translated into a molecular mechanism, an arginine side chain flip provides a charge balance. The acute sensitivity of FutA to specific radiation damage illustrates the requirement for dose limiting data collection regimes. We have used photoreduction as an advantage to study the transition of Fe(III) to Fe(II) binding state. The X-ray pump probe approach demonstrated here has the potential to become a straightforward-to-implement approach to induce redox state changes probing structural transitions. We envisage that more complex experiments could generate photoreduced states akin to anaerobic conditions that are amenable for further modification by ligand addition.

## Materials and Methods

The sections *molecular biology*; *protein purification*; *protein crystallization*; *sample preparation for serial crystallography; crystallographic data processing; structure determination and refinement*; *in crystallo UV-vis spectroscopy* are found in SI. All studies (except EPR) were performed at ambient (room) temperature. Crystallization used the natural pH of the purification buffer (0.1 M Tris buffered at pH 9.0, containing 320 mM NaCl), and 12% (w/v) PEG3350 / 0.2 M NaSCN in vapor diffusion for the home source and in batch for neutron diffraction structures. Seeded batch crystallization with 20% (w/v) PEG3350 / 0.2 M NaSCN was used for serial crystallography. Diffraction-weighted doses (DWD) reported include photoelectron escape calculation with *RADDOSE-3D* (version 2.1) (44) (for a critical discussion on dose calculation see SI).

### Home source crystal structure

Data were collected from a single crystal grown from batch crystallization and measuring 0.23 x 0.24 x 0.12 mm^3^, mounted in a 0.7 mm sealed quartz capillary on a Rigaku 007 HF (High Flux) diffractometer equipped with a HyPix 6000HE detector. The X-ray beam with a flux of 2.5 x 10^9^ ph/s at 8.1 keV was collimated at 200 µm^2^. The total exposure time of 1 hr equated to a total dose of 110 kGy.

### Neutron crystallography

For hydrogen-deuterium exchange, Fe(III) loaded FutA crystals grown from batch crystallization were transferred into a deuterated solution of the same crystallization conditions, in two subsequent exchanges for 24 hrs. Crystals with a volume larger than 0.2 mm^3^ were mounted in 1 mm sealed quartz capillaries. Data collection at BIODIFF (45), Forschungsreaktor München II (Germany) used a monochromatic neutron beam. The final dataset was merged from two isomorphous crystals collected at wavelengths of 3.1 Å (calibrated to 4DP with an Yttrium Iron Garnet powder sample).

### Serial synchrotron crystallography (SSX)

SSX data were collected at beamline I24, Diamond Light Source, using silicon chips with 12 µm apertures. For each dose series, ten images (10 ms per image) were collected at each aperture. Images were separated into individual dose points for processing to obtain ten dose points (40). Datasets above a total dose of 110 kGy were no longer isomorphous with the lowest dose point, with increased B-factors corroborating global damage.

### Serial femtosecond crystallography (SFX)

SFX data were collected at SACLA beamline BL2 EH3, Japan, using the MPCCD detector. The XFEL was operated at an X-ray energy of 11.0 keV with a pulse length of 10 fs and a repetition rate of 30 Hz. Synchronizing chip translation with the XFEL pulse, data collection took roughly 14 mins per chip.

### SFX X-ray pump probe

For the XRPP experiments, a flipper-attenuator was used to reduce the flux of alternate XFEL pulses. A fast, self-restoring rotary shutter (Branstrom Instruments, USA) mounted upstream of the sample and containing Sapphire wafer in a range of thicknesses was triggered, via TTL from a signal generator, to move the wafer into and out of the X ray beam path. Pump and probe diffraction images were separated based on total scattering intensity using the dxtbx.radial_average function from the DIALS software package (**Fig. S2**).

### UV-vis absorption spectroscopy

In solution spectra were collected in purification buffer (0.1 M Tris buffered at pH 9.0, containing 320 mM NaCl) on a Shimadzu UV-2600 spectrophotometer at a protein concentration of 4.75 mg/ml (0.14 mM). In the chemical reaction experiment, Na2S2O4 was added to a final concentration of 1.4 mM under aerobic conditions. *In crystallo* X-ray dose dependent UV-vis absorption spectroscopy was performed at ESRF beamline BM07-FIP2 with a 200 x 200 µm^2^ X-ray top-hat beam at 12.66 keV (4.1 and 5.0 x 10^11^ ph/s photon flux). Spectra were acquired at 0.4 Hz with a loop-mount crystal using a humidity controller (HC-Lab, Arinax) (46) bathed in the X-ray beam on an online microspectrophotometer with a focal volume of 50 x 50 x ∼100 µm^3^ (37, 47).

### Electron paramagnetic resonance

FutA at a concentration of 50 µM was shock-frozen in liquid nitrogen. In the chemical reduction experiment, Na2S2O4 was added to a final concentration of 500 µM under aerobic conditions prior to freezing. Data collection was carried out in EPR quartz tubes at liquid helium temperature. X-band continuous wave EPR spectra (10 Gauss modulation amplitude, 2 mW microwave power) were recorded on a Bruker eleXsys E500 spectrometer using a standard rectangular Bruker EPR cavity (ER4102T) equipped with an Oxford helium cryostat (ESR900) at 5 – 6 K.

## Acknowledgments

We thank Chris Holes for macromolecular crystallization, Peter Horton for diffraction, and Peter Roach for critical discussion at the University of Southampton (UoS). Financial support: Japan Partnering Award, Biological Sciences Research Council (BBSRC) BB/R021015/1, BB/W001950/1 to JW, MH, RO; Diamond Doctoral Studentship Programme to RB, JS, MR, CC; South Coast Biosciences Doctoral Training Partnership SoCoBio DTP BBSRC BB/T008768/1 to JS; PhD studentships by Hamburg University and the European Synchrotron Radiation Facility (ESRF) to NC, the Collaborative Computing Project 4 (CCP4) to LC (#7920S22020007); the Institute for Life Sciences (Southampton) to CC; Wellcome Investigator Award 210734/Z/18/Z to AMO; Royal Society Wolfson Fellowship RSWF\R2\182017 to AMO. We acknowledge facility access to the National Crystallography Service (NCS) Southampton; DLS MX15722, NT14493, NT23570; SACLA 2022A8002, 2022B8041; Forschungsreaktor München ID:16106; ESRF BM07-FIP2 and *ic*OS, MX2373, MX2374; Diamond Light Source (DLS) UK XFEL hub and ESRF for travel support. We are grateful to beamline staff, in particular Shigeki Owada and Kensuke Tono at SACLA.

## References

1. K. Hans Wedepohl, The composition of the continental crust. Geochimica et Cosmochimica Acta 59, 1217–1232 (1995).

2. P. W. Boyd et al., Mesoscale Iron Enrichment Experiments 1993-2005: Synthesis and Future Directions. Science 315, 612–617 (2007).

3. W. Stumm, B. Sulzberger, The cycling of iron in natural environments: Considerations based on laboratory studies of heterogeneous redox processes. Geochimica et Cosmochimica Acta 56, 3233–3257 (1992).

4. M. L. Wells, N. M. Price, K. W. Bruland, Iron chemistry in seawater and its relationship to phytoplankton: a workshop report. Marine Chemistry 48, 157–182 (1995).

5. H. W. Rich, F. M. M. Morel, Availability of well-defined iron colloids to the marine diatom Thalassiosira weissflogii. Limnology and Oceanography 35, 652–662 (1990).

6. J. A. Raven, M. C. W. Evans, R. E. Korb, The role of trace metals in photosynthetic electron transport in O2-evolving organisms. Photosynthesis Research 60, 111–150 (1999).

7. S. Richier et al., Abundances of Iron-Binding Photosynthetic and Nitrogen-Fixing Proteins of Trichodesmium Both in Culture and In Situ from the North Atlantic. PLoS ONE 7, e35571 (2012).

8. J. T. Snow et al., Quantifying Integrated Proteomic Responses to Iron Stress in the Globally Important Marine Diazotroph Trichodesmium. PLOS ONE 10, e0142626 (2015).

9. Z. S. Kolber et al., Iron limitation of phytoplankton photosynthesis in the equatorial Pacific Ocean. Nature 371, 145–149 (1994).

10. M. C. Moore et al., Large-scale distribution of Atlantic nitrogen fixation controlled by iron availability. Nature Geoscience 2, 867–871 (2009).

11. M. A. Huston, S. Wolverton, The global distribution of net primary production: resolving the paradox. Ecological Monographs 79, 343–377 (2009).

12. P. Flombaum et al., Present and future global distributions of the marine cyanobacteria Prochlorococcus and Synechococcus. Proceedings of the National Academy of Sciences 110, 9824–9829 (2013).

13. S. J. Biller, P. M. Berube, D. Lindell, S. W. Chisholm, Prochlorococcus: the structure and function of collective diversity. Nature Reviews Microbiology 13, 13–27 (2015).

14. F. Partensky, W. R. Hess, D. Vaulot, Prochlorococcus, a Marine Photosynthetic Prokaryote of Global Significance. Microbiology and Molecular Biology Reviews 63, 106–127 (1999).

15. P. M. Berube et al., Single cell genomes of Prochlorococcus, Synechococcus, and sympatric microbes from diverse marine environments. Sci Data 5, 180154 (2018).

16. R. Sutak, J.-M. Camadro, E. Lesuisse, Iron Uptake Mechanisms in Marine Phytoplankton. Frontiers in Microbiology 11 ((2020)).

17. M. Sandy, A. Butler, Microbial Iron Acquisition: Marine and Terrestrial Siderophores. Chemical Reviews 109, 4580–4595 (2009).

18. D. B. Rusch, A. C. Martiny, C. L. Dupont, A. L. Halpern, J. C. Venter, Characterization of Prochlorococcus clades from iron-depleted oceanic regions. Proceedings of the National Academy of Sciences 107, 16184–16189 (2010).

19. R. R. Malmstrom et al., Ecology of uncultured Prochlorococcus clades revealed through single-cell genomics and biogeographic analysis. International Society for Microbial Ecology Journal 7, 184–198 (2013).

20. G. Rocap et al., Genome divergence in two Prochlorococcus ecotypes reflects oceanic niche differentiation. Nature 424, 1042–1047 (2003).

21. H. Katoh, N. Hagino, A. R. Grossman, T. Ogawa, Genes essential to iron transport in the cyanobacterium Synechocystis sp. strain PCC 6803. Journal of Bacteriology 183, 2779–2784 (2001).

22. A. Badarau et al., FutA2 is a ferric binding protein from Synechocystis PCC 6803. Journal of Biological Chemistry 283, 12520–12527 (2007).

23. P. Exss-Sonne, J. TÖlle, K. P. Bader, E. K. Pistorius, K.-P. Michel, The IdiA protein of Synechococcus sp. PCC 7942 functions in protecting the acceptor side of Photosystem II under oxidative stress. Photosynthesis Research 63, 145–157 (2000).

24. J. TÖlle et al., Localization and function of the IdiA homologue Slr1295 in the cyanobacterium Synechocystis sp. strain PCC 6803. Microbiology 148, 3293–3305 (2002).

25. K. P. Michel, E. K. Pistorius, Adaptation of the photosynthetic electron transport chain in cyanobacteria to iron deficiency: The function of IdiA and IsiA. Physiologia Plantarum 120, 36–50 (2004).

26. H. Katoh, N. Hagino, T. Ogawa, Iron-binding activity of FutA1 subunit of an ABC-type iron transporter in the cyanobacterium Synechocystis sp. Strain PCC 6803. Plant and Cell Physiology 42, 823–827 (2001).

27. D. Polyviou et al., Structural and functional characterization of IdiA/FutA (Tery_3377), an iron-binding protein from the ocean diazotroph Trichodesmium erythraeum. Journal of Biological Chemistry 293, 18099–18109 (2018).

28. E. F. Garman, M. Weik, Radiation Damage in Macromolecular Crystallography. Methods Mol Biol 1607, 467–489 (2017).

29. J. A. R. Worrall, M. A. Hough, Serial femtosecond crystallography approaches to understanding catalysis in iron enzymes. Curr Opin Struc Biol 77 ((2022)).

30. V. Pfanzagl et al., X-ray–induced photoreduction of heme metal centers rapidly induces active-site perturbations in a protein-independent manner. Journal of Biological Chemistry 295, 13488–13501 (2020).

31. S. Horrell et al., Fixed Target Serial Data Collection at Diamond Light Source. J Vis Exp 10.3791/62200 (2021).

32. S. Wang et al., A novel mode of ferric ion coordination by the periplasmic ferric ion-binding subunit FbpA of an ABC-type iron transporter from Thermus thermophilus HB8. Acta Crystallographica Section D 70, 196–202 (2014).

33. G. Palmer, The electron paramagnetic resonance of metalloproteins. Biochem Soc Trans 13, 548–560 (1985).

34. H. N. Chapman, X-Ray Free-Electron Lasers for the Structure and Dynamics of Macromolecules. Annu Rev Biochem 88, 35–58 (2019).

35. K. Nass et al., Structural dynamics in proteins induced by and probed with X-ray free-electron laser pulses. Nat Commun 11, 1814 (2020).

36. J. H. Beale et al., Successful sample preparation for serial crystallography experiments. Journal of Applied Crystallography 52, 1385–1396 (2019).

37. D. von Stetten et al., In crystallo optical spectroscopy (icOS) as a complementary tool on the macromolecular crystallography beamlines of the ESRF. Acta Crystallogr D Biol Crystallogr 71, 15–26 (2015).

38. A. M. Orville, N. Elango, J. D. Lipscomb, D. H. Ohlendorf, Structures of competitive inhibitor complexes of protocatechuate 3,4-dioxygenase: multiple exogenous ligand binding orientations within the active site. Biochemistry 36, 10039–10051 (1997).

39. M. A. Hough, R. L. Owen, Serial synchrotron and XFEL crystallography for studies of metalloprotein catalysis. Curr Opin Struct Biol 71, 232–238 (2021).

40. A. Ebrahim et al., Dose-resolved serial synchrotron and XFEL structures of radiation-sensitive metalloproteins. International Union of Crystallography Journal 6, 543–551 (2019).

41. I. G. Denisov, D. C. Victoria, S. G. Sligar, Cryoradiolytic reduction of heme proteins: Maximizing dose-dependent yield. Radiation Physics and Chemistry 76, 714–721 (2007).

42. J. S. Fraser et al., Accessing protein conformational ensembles using room-temperature X-ray crystallography. Proc Natl Acad Sci U S A 108, 16247–16252 (2011).

43. S. L. S. Storm et al., Measuring energy-dependent photoelectron escape in microcrystals. IUCrJ 7, 129–135 (2020).

44. C. S. Bury, J. C. Brooks-Bartlett, S. P. Walsh, E. F. Garman, Estimate your dose: RADDOSE-3D. Protein Science 27, 217–228 (2018).

45. T. S. A. Ostermann, BIODIFF: Diffractometer for large unit cells. Journal of large-scale research facilities 1, A2 (2015).

46. J. Sanchez-Weatherby et al., Improving diffraction by humidity control: a novel device compatible with X-ray beamlines. Acta Crystallogr D Biol Crystallogr 65, 1237–1246 (2009).

47. J. McGeehan et al., Colouring cryo-cooled crystals: online microspectrophotometry. J Synchrotron Radiat 16, 163–172 (2009).

