## Supplementary Information for "A redox switch allows binding of Fe(II) and Fe(III) ions in the cyanobacterial iron binding protein FutA from *Prochlorococcus*"

PNAS

**Rachel Bolton *et al.***

**This supplement includes:**

Extended Materials and Methods
Supplementary Text
Tables S1 to S4
Figures S1 to S5

**Extended Materials and Methods**

*Molecular biology. Prochlorococcus* MED4 *futA* was cloned into pET-24b(+) using the NdeI / HindIII restriction sites, excluding the region encoding the N-terminal signal peptide, amino acids 27-340 (UniProt ID: Q7V0T9) as predicted by *SignalP* (1).

*Protein purification.* Transformed *Escherichia coli* BL21 (DE3) cells (NEB) were cultured in 3 L baffled flasks in 1 L lysogeny broth containing 50 $\mu$g ml$^{-1}$ kanamycin, and incubated in a shaker at 130 RPM, 37 °C. The temperature was reduced to 18 °C when the cell culture reached an $OD_{600}$ of 0.4. Protein expression was induced at an $OD_{600}$ of ~0.6 by addition of IPTG (final concentration 1 mM). Cells were harvested after 20 hrs by centrifugation at 4000 x g (Avanti Jxn-26, JLA-8.1000 rotor). Cell pellets (2-4g) were resuspended in 25 ml IBB buffer (0.1 M Tris buffered at pH 9, containing 0.5 M NaCl, 1% Triton-X, 5 mM $MgCl_2$ and 10 mM β-mercaptoethanol). For lysis 2 mg ml$^{-1}$ lysozyme was added, and cells were left for 30 min before sonication for total pulse time of 150 seconds (Q700 Sonicator, 10 second pulse duration with 20 seconds between pulses). Inclusion bodies were harvested by centrifugation (40 mins, 125 000 x g, 4 °C, Optima XPN-80, Type 70 Ti rotor). The pellet was washed in IBB containing 2 M urea, followed by centrifugation (as above). Solubilization was carried out by incubation in 200 mM Tris buffered at pH 9, containing 6 M urea, 10 mM β-mercaptoethanol (1 hrs, 4 °C). After removing cellular debris by centrifugation (as above), a rapid dilution protocol was carried out to refold the protein. The salute was loaded into a syringe with a fine needle and slowly added directly into 2 L of stirring 0.2 M Tris buffered at pH 9.0, containing 0.2 M NaCl, 0.4 M L-Arginine and 0.1 mM $NH_4Fe(SO_4)_2$. After incubation at 4 °C for 48 h, the refolding buffer was concentrated to 150 ml using an Amicon Stirred Cell (10,000 Da Ultrafiltration Disk, Merck). Dialysis against 2 L 100 mM Tris buffered at pH 9.0, containing 145 mM NaCl for 24 hrs at 4 °C was followed by capture on a 5 ml HiTrap SP XL column (GE Healthcare) at room temperature. Step-elution with 0.1 M Tris buffered at pH 9.0, containing 320 mM NaCl was followed by size-exclusion chromatography on a HiLoad 16/60 Superdex 200 column (GE Healthcare) using 50 mM Tris buffered at pH 9.0, containing 300 mM NaCl at room temperature. Fractions containing monomeric FutA were pooled and concentrated using a Vivaspin 20 Centrifugal Concentrator, MWCO 10,000 Da (Sartorius) at 4 °C and stored at 4 °C or room temperature.

*Protein Crystallization.* FutA at a concentration of ~50 mg ml$^{-1}$ was crystallized at room temperture at the final pH from purification (pH 9.0). The specific crystallization conditions used for each dataset described are described in the main manuscript, all crystals burgundy red. FutA typically crystallized at a range of precipitant concentrations. For vapor diffusion crystallization, 1 $\mu$l protein was mixed with 1 $\mu$l 0.2 M sodium thiocyanate containing 10 - 35 % (w/v) PEG 3350 and set up in 24-well XRL plates (Molecular Dimensions). Crystals with 10 – 200 $\mu$m in the longest dimension appeared within 1 day. For the home source and neutron diffraction experiments, batch crystallization was used where 10 $\mu$l of protein was mixed with 10 $\mu$l of 0.2 M sodium thiocyanate containing 12 % (w/v) PEG 3350 in a microcentrifuge tube. Crystals with 200 – 1500 $\mu$m in the longest dimension appeared within 3 days. For serial crystallography, seeded batch crystallization was used where seeds were generated by mixing 10 $\mu$l of FutA crystals obtained from vapor diffusion droplets with 40 $\mu$l 20% PEG 3350, followed by vortexing with Seed Bead for 180 s (Hampton Research). Seed stock aliquots (5 $\mu$l) were shock frozen and diluted 1:100 with 0.2 M sodium thiocyanate, 20 % (w/v) PEG 3350 prior to use. For crystallization, 50 $\mu$l protein was mixed with 75 $\mu$l diluted seed stock and 75 $\mu$l of 0.2 M sodium thiocyanate containing 20 % (w/v) PEG 3350. Crystals with 10 – 20 $\mu$m in the longest dimension appeared within 30 minutes as described previously (2).

*Sample preparation for serial crystallography.* Optimization of crystallization for serial crystallography was described previously (2). We used a fixed-target silicon chip, each accommodating 25,600 apertures, to deliver microcrystals to the X-ray interaction region (3). The crystal slurry (typically 150 $\mu$l) was loaded onto a glow-discharged chip containing 7 or 12 $\mu$m sized apertures within a humidity-controlled chamber, collected by applying vacuum, and sealing the chip between two sheets of 6 μm thick Mylar. Crystal slurries had to be prepared directly before the experiment to avoid crystal ageing that manifested as loss of diffraction.

*Crystallographic data processing, structure determination and refinement.* Data collection for all datasets was performed at ambient (room) temperature, with data collection and refinement statistics as given in **Table S1**. The SFX 5 kGy and XRPP 320 kGy datasets show higher B-values than the other three datasets, likely rooted in de/hydration effects of the fixed target chip mount and through incident X-ray irradiation, in particular in the XRPP experiment. The ambient temperature where the data was collected is reported, and with 25 °C was higher for the XFEL experiment. The home source diffraction data were integrated with *XDS* (4), and scaled / merged using *POINTLESS* and *AIMLESS* (5). The Neutron diffraction data were integrated using *HKL2000* (6) and scaled / merged using *SCALEPACK* (6). The diffraction data for the SSX dose series was indexed and integrated using *dials.stills_process* (*DIALS* v2.0) (7) and scaled using *cctbx.prime* (8). Results for two dose-series are shown in **Table S2**, using 5 kGy and 22 kGy dose-slicing. B-factor sharpening was applied in scaling to correct for the increase in B-factors in each series. *SCALEit* (9) was used to derive the isomorphous difference between two datasets ($D_{iso}$), and scale the differences in observed structure factors between datasets. SFX diffraction data were stored in a hdf5 stream, applying indexing and pre-filtering for diffraction hits with *Cheetah* (10). Diffraction hits were indexed and integrated with *dials.stills_process* (*DIALS* v3.0) (7). An image mask was generated manually using *dials.image_viewer* to remove the beam stop shadow and monocrystalline Si diffraction spots arising from the chips. Integrated patterns were scaled and merged using the *DIALS* module *cctbx.xfel.merge* (11). Molecular replacement with *MOLREP* (12) used the *Synechocystis* PCC 6803 FutA2 as search model (PDB: 2PT1). *COOT* (13), and *REFMAC5* (14) were used for iterative model building, refinement and validation. Coordinates and structure factors were deposited with the PDB under accession numbers 8OEM (Home Source), 8OEN (Neutron), 8C4Y (SFX), 8OGG (SSX 5 kGy), and 8OEI (SFX 350kGy), using the EBI validation suite.

*In crystallo UV-vis spectroscopy.* Online micro spectrophotometry at beamline ESRF BM07-FIP2 used 400 μm optical fibre to connect a balanced deuterium-halogen lamp (Mikropack DH2000-BAL, Ocean Optics) to the higher objective, while a 600 μm optical fibre connected the lower objective to a fixed-grating spectrophotometer equipped with a CCD detector (QE65 Pro, Ocean Optics). Spectra were acquired at 0.4 Hz (250 ms acquisition time averaged 10 times) on several crystals with a volume between 160 x 70 x 50 $\mu m^3$ and 200 x 90 x 80 $\mu m^3$. Crystals were oriented to optimize the signal-to-noise ratio of the spectra and maintained still during X-ray exposure in a loop-mount using a humidity controller (HC-Lab, Arinax) (15). Data analysis was performed using a suite of in-house Python scripts (https://github.com/ncara/icOS) with the *NumPy* (16), *pandas* (17), *SciPy* (18) and *Matplotlib* (19) packages. Spectra were first baseline-corrected by subtraction of a constant corresponding to the average absorption between 800 and 880 nm. They were then smoothed using a Savitzky-Golay filter (parameters: $3^{rd}$ order polynom; 25-data point smoothing window).

**Supplementary Text**

**Comparison of the iron coordination in the NMX and SFX structures.**

Further investigation of the iron binding site was carried out by performing refinement runs of the NMX and SFX models using REFMAC5 (20). The latest procedure implemented in REFMAC5 available in CCP4 8.0 includes the refinement of structures obtained by NMX (manuscript submitted, (21)).

Crystallographic refinement was carried out using two alternative strategies. In a first approach, refinement was performed without restraints between the iron center and its coordination ligands. In a second approach, restraints were employed between the metal center and its four coordinating tyrosine residues (Tyr13, Tyr143, Tyr199, Tyr200). Restraint values were derived from the Crystallography Open Database (COD) (22) and two Fe-O target values (1.800 Å and

Fe-O = 2.004 Å) were tested. In the latter approach we also utilized four different weights for the Fe-O restrains with sigma values ranging from 0.01 Å (tight) to 0.04 Å (soft). Coordination distances following refinement are reported in **Table S2**.

It appears that whilst Fe-ligand distances in the SFX structure are rather insensitive to the restraints and sigma values employed, those for the NMX model exhibit a behavior that is more biased toward the chosen target value. The average coordination distance in the NMX model appears to be marginally larger than that for the SFX model, with values of 2.10 Å and 1.93 Å, respectively. In general, differences in apparent distances are to be expected as NMX reports on the position of the atomic nuclei whilst X-ray methods report on the electron distribution. We note, however, that the Fe-water distance in the NMX model is ~0.5 Å larger than that in the SFX model. Additionally, the neutron structure features a water molecule in H-bonding distance to the iron coordinating water that is not seen in the SFX model. Overall, under the reasonable assumption that both structures represent the same oxidation state for the iron center, it would appear that a limited degree of coordination flexibility is possible.

**Discussion of dose calculation**

Raddose-3D was used in all cases for comparability, this being the best available tool to estimate dose (23). For SFX XRPP, RADDOSE-3D rather than RADDOSE-XFEL (24) was used as we were probing the effects of the total dose deposited by the pump pulse 33 milliseconds after the pulse rather than the time resolved evolution of dose within the duration of the pulse. Diffraction-weighted doses (DWD) with inclusion of photoelectron escape are reported in all cases since it is a common metric to compare data from different sources.

In this communication we describe differences in metal coordination based on dose. However, we accept that the different regimen described in the experiments could lead to the different observations. In the case of the SFX experiment, the beam was small with respect to the crystal, leading to the x-ray exposed region being surrounded by a large nonexposed crystal volume. In the SSX experiment the crystal is bathed in the beam (at least in two dimensions, as FutA crystals have an axis ratio of typ. 2.5:1:1). X-ray exposure at an XFEL differs from the more conventional SSX experiment by twelve orders of magnitude. The home source experiment did not only differ in dose rate and duration of the full experiment, but also in beam and crystal size. The difference in the behavior of the iron coordinating water between the home source, SSX, and SFX XRPP experiments could be attributed to differences in hygroscopic stability of the sample delivery systems (quartz capillary vs mylar sealed chip) or possible dehydration differences between the individual crystal preparations.

The mechanisms by which redox chemistry plays out in the XRPP experiment after the first 10fs XFEL pulse may well be rather different to a conventional experiment at a synchrotron source with a constant X-ray exposure. Generally, it is the production of solvated photoelectrons from radiolysis of water molecules within the crystal that primarily drive dose-dependent reduction of redox centres. The cascade of reactions leading to production of these photoelectrons however may well be different in these two scenarios. While under cryogenic conditions radiation damage has been shown to be (essentially) purely dose dependent, at room temperature there are clearly time-dependent processes occurring as a result of X-ray exposure, as well as dose driven processes.

For the optical spectroscopic data, we also used DWD to allow comparableness, but we note that a spectroscopic experiment performed on a crystallography beamline with a top-hat-shaped beam irradiating a still crystal would rather use the ADER metric (Average Dose over the Exposed Region), since doses corresponding to successive spectra increase linearly with exposure time. In contrast, the DWD of an oscillation experiment takes into account that the first images "see" fewer X-rays than the ones at the end of the experiment (24). Calculated as ADER, reported doses would approximately double for the spectroscopic experiment with a top-hat-shaped X-ray beam. It was previously discussed that ADER might be used when microcrystals and microbeams are used (25), though it is presently unclear how the XRPP data should be treated.

For the above reasons, interpretation of the data on the dose variable alone may not be sufficient to explain the observations made. It is evident that further study on the effects of XFEL pulses *versus* those of continuous synchrotron exposure on the dose and its structural consequences is required.

**Analysis of sequence conservation in the iron binding site of FutA proteins**

The data in this manuscript suggest that *Prochlorococcus* MED4 FutA can bind iron in the Fe(**III**) and in the Fe(**II**) redox states, using an arginine switch mechanism. This observation raises the question whether this concept can be extended to other FutA homologues. We therefore carried out multiple sequence alignment analysis (MSA) across a set of known FutA homologues, **Fig. S5**.

The bacterial species selected here are gram-negative, free-living marine bacteria that are found in oligotrophic ocean waters. Of these, *Prochlorococcus* MED4 and *Synechocystis* PCC 6803 are capable of carbon fixation, whereas *T. erythraeum* and *C. chwakensis* are also capable of nitrogen fixation. *T. thermophilus* is a heterotrophic, extremophile, isolated from deep-sea thermal vents. Species with a single FutA homologue in their genome were *Prochlorococcus* MED4 (UniProt ID: Q7V0T9), *Trichodesmium erythraeum* (UniProt ID: Q10Z45), *Crocosphaera chwakensis* (A3IPT8), and *Thermus thermophilus* (UniProt ID: Q5SHV2). In contrast, *Synechocystis* PCC 6803 has two FutA homologies, denoted as FutA1 (UniProt ID: P72827) and FutA2 (UniProt ID: Q55835).

The sequences of the FutA homologues were aligned with Clustal Omega (26) and visualized with JalView (27), as shown in **Fig. S5**. Conservation of the physico-chemical properties of the amino acids across the sequence alignment is shown as a gradient from blue (low conservation) to yellow (high conservation) (27). The N-terminal signal peptide of *Prochlorococcus* MED4 (amino acids 1-27) as predicted by *SignalP* (1) (see *Molecular biology*, *Extended Materials and Methods*) was excluded from the alignment.

For species encoding a single FutA protein the arginine residue equivalent to Arg203 in *Prochlorococcus* MED4 FutA was conserved. Interestingly, this amino acid is even conserved *Synechocystis* PCC 6803 which has two FutA homologues, which may suggest that FutA1 and FutA2 both have capacity to bind Fe(**III**) or Fe(**II**) iron. This observation aligns with gene knockout studies conducted by us (28) demonstrating a degree of redundancy between the two FutA homologues.

The iron binding site in FpbA from *T. thermophilus* differs from the other proteins, as the residues equivalent to *Prochlorococcus* MED4 FutA His12 and Tyr13 are exchanged to Glycine and Glutamine, respectively. Despite this, the residue equivalent to Arg203 is conserved (Arg223). As FbpA switches from an open (PDB: 3WAE) to a closed conformation (PDB: 4ELR) upon iron binding, a carbonate ion is lost from the binding site and Arg223 is repositioned away from the binding site, **Fig. S4** (29, 30). This ability of Arg223 to act as a structural switch and maintain a net neutral charge in the binding site is therefore highly similar to the observed structural rearrangement of Arg203 in *Prochlorococcus* MED4 FutA.

**Table S1.** Data collection and refinement statistics for FutA structures reported (space group $P2_1$)

| | Home Source | Neutron | SFX | SSX 5 kGy | SFX 350kGy |
|---|---|---|---|---|---|
| Temperature (°C) | 21 | 21 | 25 | 21 | 25 |
| Wavelength | 1.54 | 3.10 | 1.13 | 0.97 | 1.13 |
| # Integrated Lattices | | | 78,743 | 5,278 | 24,378 |
| # Merged Lattices | | | 77,936 | 5,170 | 24,141 |
| Unit Cell (a, b, c; Å) | 39.4, 78.0, 48.0 | 39.5, 78.3, 47.9 | 39.1, 78.3, 47.4 | 39.7, 78.7, 48.4 | 39.4, 78.2, 48.0 |
| β angle (°) | 98.2 | 97.4 | 97.4 | 97.8 | 97.9 |
| Resolution (all, Å) | 47.50 – 1.70 | 24.95 – 2.1 | 30.10 – 1.60 | 40.97 – 1.76 | 32.42 – 1.65 |
| Resolution (HR, Å) | 1.73 – 1.70 | 2.18 – 2.1 | 1.63 – 1.60 | 1.79 – 1.76 | 1.68 – 1.65 |
| $R_{split}$ / $R_{pim}$[1] | 0.009 (0.089) | 0.098 (0.332) | 0.053 (0.089) | 0.235 (0.684) | 0.134 (0.691) |
| CC ½ (%)[1] | 100.0 (98.6) | 97.1 (68.1) | 99.6 (90.6) | 92.3 (43.1) | 97.0 (40.0) |
| $I/\sigma I$[1] | 71.7 (11.6) | 4.8 (2.0) | 12.1 (4.6) | 3.14 (0.41) | 3.67 (0.43) |
| Completeness (%)[1] | 97.9 (83.5) | 81.0 (52.6) | 99.8 (100.0) | 100.0 (100.0) | 100.0 (100.0) |
| Multiplicity[1] | 65.4 (42.1) | 1.9 (1.1) | 618.7 (274.5) | 26.2 (19.1) | 165.2 (82.4) |
| Unique Reflections[1] | 30,881 (1,369) | 13,731 (890) | 37,266 (1,877) | 29,256 (1,467) | 34,653 (1676) |
| Wilson B-factor (Å$^2$) | 14.10 | 10.07 | 12.09 | 24.28 | 20.98 |
| | **Home Source yellow** | **Neutron blue** | **SFX green** | **SSX 5 kGy purple** | **SFX 350 kGy green** |
| PDB Code | 8OEM | 8OEN | 8C4Y | 8OGG | 8OEI |
| Resolution (Å) | 47.50 – 1.70 | 24.95 – 2.1 | 30.10 – 1.60 | 40.97 – 1.76 | 32.42 – 1.65 |
| Rwork/Rfree | 0.153 / 0.180 | 0.182 / 0.250 | 0.190 / 0.208 | 0.202 / 0.241 | 0.162 / 0.189 |
| # Reflections all/free | 30847 / 1,588 | 13,731 / 698 | 37,266 /1,919 | 29256 / 1,503 | 34,653 / 1,786 |
| Number of Atoms | | | | | |
| Protein | 2558 | 4984 | 2485 | 2509 | 2572 |
| Ion | 1 | 1 | 1 | 1 | 1 |
| Water | 117 | 81 | 86 | 81 | 105 |
| Clashscore (all)[2] | 2.31 | 3.41 | 1.79 | 1.58 | 2.51 |
| Ramachandran | | | | | |
| Preferred | 305 | 298 | 304 | 301 | 302 |
| Allowed | 4 | 11 | 5 | 8 | 7 |
| Outliers | 1 | 1 | 1 | 1 | 1 |
| Z-score[2] | 0.20±0.40 | -0.20±0.43 | 0.07±0.41 | -0.69±0.41 | 0.17±0.42 |
| B-factors (Å$^2$) | | | | | |
| Protein | 19.71 | 22.31 | 17.84 | 33.32 | 27.06 |
| Water | 25.32 | 13.41 | 23.32 | 48.19 | 33.05 |
| R.M.S Deviations | | | | | |
| Bond Lengths (Å) | 0.013 | 0.005 | 0.013 | 0.006 | 0.009 |
| Bad bonds[2] | 1 / 2,612 | 0 / 2,506 | 1 / 2,530 | 0 / 2,557 | 0 / 2,625 |
| Bond Angles (°) | 1.88 | 1.14 | 1.76 | 1.41 | 1.54 |
| Bad angles[2] | 8 / 3,537 | 0 / 3,377 | 5 / 3,416 | 1 / 3,456 | 3 / 3,553 |
| Molprobity score | 1.01 | 1.13 | 0.94 | 1.02 | 1.03 |

[1]High resolution statistics in parentheses; [2]As determined by MolProbity (31)

**Table S2.** Bond lengths for the Fe coordination cage in the SFX and NMX structures following different refinement strategies as indicated in the text. All tabulated bond lengths are in Å.

| | no link | Fe-Tyr link (1.8 Å) | | | | Fe-Tyr link (2.004 Å) | | | |
|---|---|---|---|---|---|---|---|---|---|
| SFX | | | | | | | | | |
| Fe-O | | 0.01 $\sigma$ | 0.02 $\sigma$ | 0.03 $\sigma$ | 0.04 $\sigma$ | 0.01 $\sigma$ | 0.02 $\sigma$ | 0.03 $\sigma$ | 0.04 $\sigma$ |
| Fe-Tyr13 | 1.80 | 1.80 | 1.80 | 1.80 | 1.80 | 1.94 | 1.86 | 1.83 | 1.82 |
| Fe-Tyr143 | 1.97 | 1.87 | 1.93 | 1.95 | 1.96 | 1.99 | 1.98 | 1.98 | 1.98 |
| Fe-Tyr199 | 1.87 | 1.83 | 1.85 | 1.86 | 1.87 | 1.95 | 1.91 | 1.89 | 1.89 |
| Fe-Tyr200 | 1.82 | 1.81 | 1.82 | 1.82 | 1.82 | 1.93 | 1.87 | 1.84 | 1.83 |
| Fe-Water | 2.17 | 2.16 | 2.16 | 2.16 | 2.16 | 2.16 | 2.16 | 2.16 | 2.16 |
| Average | 1.93 | 1.89 | 1.91 | 1.92 | 1.92 | 1.99 | 1.96 | 1.94 | 1.94 |
| NMX | | | | | | | | | |
| Fe-Tyr13 | 2.09 | 1.81 | 1.85 | 1.90 | 1.95 | 2.01 | 2.02 | 2.04 | 2.05 |
| Fe-Tyr143 | 1.95 | 1.81 | 1.83 | 1.85 | 1.87 | 2.0 | 2.0 | 1.99 | 1.99 |
| Fe-Tyr199 | 1.83 | 1.80 | 1.80 | 1.80 | 1.81 | 2.0 | 1.98 | 1.96 | 1.94 |
| Fe-Tyr200 | 1.91 | 1.81 | 1.83 | 1.85 | 1.87 | 2.0 | 1.99 | 1.98 | 1.97 |
| Fe-Water | 2.72 | 2.64 | 2.66 | 2.68 | 2.69 | 2.60 | 2.61 | 2.63 | 2.64 |
| Average | 2.10 | 1.97 | 1.99 | 2.02 | 2.04 | 2.12 | 2.12 | 2.12 | 2.12 |

**Table S3**. Full data collection and refinement statistics for two SSX dose series reported in space group $P2_1$ Data collection was carried out at 21 °C at an energy of X-ray Energy of 12.8 keV for 10 consecutive exposures to give dose points at 5 kGy interval.

Data collection statistics for the 5 kGy SSX dose-series

| **Data Collection** | **5 kGy** | **10 kGy** | **15 kGy** | **20 kGy** | **25 kGy** |
|---|---|---|---|---|---|
| Number of Integrated Lattices | 5,278 | 5,089 | 5,197 | 5,158 | 5,227 |
| Number of Merged Lattices | 5,170 | 4,984 | 4,916 | 4,946 | 4,999 |
| Unit Cell a, b, c (Å) | 39.7, 78.7, 48.4 | 39.7, 78.7, 48.4 | 39.7, 78.7, 48.4 | 39.7, 78.7, 48.4 | 39.7, 78.7, 48.4 |
| β (°) | 97.8 | 97.8 | 97.8 | 97.8 | 97.8 |
| Resolution, overall (Å) | 40.97 – 1.76 | 40.97 – 1.76 | 40.97 – 1.76 | 40.97 – 1.76 | 40.97 – 1.76 |
| Resolution, high (Å) | 1.79 – 1.76 | 1.79 – 1.76 | 1.79 – 1.76 | 1.79 – 1.76 | 1.79 – 1.76 |
| $R_{split}$[1] | 23.5 (68.4) | 25.1 (70.8) | 20.6 (95.8) | 24.9 (71.8) | 24.3 (72.8) |
| CC ½ (%)[1] | 92.3 (43.1) | 91.0 (38.5) | 91.0 (38.9) | 89.8 (41.4) | 90.7 (42.5) |
| $I/\sigma I$[1] | 3.14 (0.41) | 3.24 (0.41) | 3.29 (0.42) | 3.33 (0.41) | 3.39 (0.40) |
| Completeness (%)[1] | 100.0 (100.0) | 100.0 (100.0) | 100.0 (100.0) | 100.0 (100.0) | 100.0 (100.0) |
| Multiplicity[1] | 26.2 (19.1) | 25.8 (18.8) | 25.7 (18.8) | 25.5 (18.6) | 26.1 (19.0) |
| Unique Reflections[1] | 29,256 (1,467) | 29,256 (1,469) | 29,257 (1472) | 29,258 (1471) | 29,258 (1476) |
| Wilson B-factor (Å$^2$) | 24.28 | 24.73 | 24.93 | 25.17 | 25.52 |
| **Data Collection** | **30 kGy** | **35 kGy** | **40 kGy** | **45 kGy** | **50 kGy** |
| Number of Integrated Lattices | 5,372 | 5,397 | 5,410 | 5,444 | 5,400 |
| Number of Merged Lattices | 5,119 | 5,110 | 5,115 | 5,160 | 5,147 |
| Unit Cell a, b, c (Å) | 39.7, 78.7, 48.4 | 39.7, 78.7, 48.4 | 39.7, 78.7, 48.4 | 39.7, 78.7, 48.4 | 39.7, 78.8, 48.4 |
| β (°) | 97.8 | 97.8 | 97.8 | 97.8 | 97.7 |
| Resolution, overall (Å) | 40.97 – 1.76 | 40.97 – 1.76 | 40.97 – 1.76 | 40.97 – 1.76 | 40.97 – 1.76 |
| Resolution, high (Å) | 1.79 – 1.76 | 1.79 – 1.76 | 1.79 – 1.76 | 1.79 – 1.76 | 1.79 – 1.76 |
| $R_{split}$[1] | 24.5 (76.4) | 24.2 (75.4) | 24.1 (74.9) | 23.6 (78.3) | 23.5 (79.1) |
| CC ½ (%)[1] | 91.3 (37.2) | 90.9 (31.1) | 91.5 (38.7) | 92.0 (34.9) | 91.4 (32.8) |
| $I/\sigma I$[1] | 3.24 (0.37) | 3.26 (0.39) | 3.31 (0.37) | 3.24 (0.36) | 3.18 (0.35) |
| Completeness (%)[1] | 100.0 (100.0) | 100.0 (100.0) | 100.0 (100.0) | 100.0 (100.0) | 100.0 (100.0) |
| Multiplicity[1] | 26.6 (19.4) | 26.9 (19.6) | 26.5 (19.4) | 26.8 (19.7) | 26.8 (19.6) |
| Unique Reflections[1] | 29,263 (1,475) | 29,265 (1,471) | 29,266 (1,476) | 29,270 (1,476) | 29,273 (1,479) |
| Wilson B-factor (Å$^2$) | 26.00 | 26.21 | 26.67 | 27.12 | 27.52 |

[1]High resolution statistics in parentheses

**Table S4**. Full data collection and refinement statistics for two SSX dose series reported in space group $P2_1$ (next pages). Data collection was carried out at 21 °C at an energy of X-ray Energy of 12.8 keV for 10 consecutive exposures to give dose points at 22 kGy interval.

Data collection statistics for the 22 kGy SSX dose-series

| **Data Collection** | **22 kGy** | **44 kGy** | **66 kGy** | **88 kGy** | **110 kGy** |
|---|---|---|---|---|---|
| Number of Integrated Lattices | 9,015 | 9,760 | 10,114 | 9,914 | 9,533 |
| Number of Merged Lattices | 8,989 | 9,723 | 10,079 | 9,878 | 9,506 |
| Unit Cell a, b, c (Å) | 39.5, 78.2, 48.1 | 39.4, 78.2, 48.1 | 39.4, 78.3, 48.0 | 39.4, 78.3, 48.0 | 39.4, 78.3, 48.0 |
| β (°) | 97.8 | 97.8 | 97.7 | 97.7 | 97.7 |
| Resolution, overall (Å) | 40.71 – 2.10 | 40.71 – 2.10 | 40.71 – 2.10 | 40.71 – 2.10 | 40.72 – 2.10 |
| Resolution, high (Å) | 2.14 – 2.10 | 2.14 – 2.10 | 2.14 – 2.10 | 2.14 – 2.10 | 2.14 – 2.10 |
| $R_{split}$[1] | 20.5 (24.7) | 18.5 (20.2) | 16.9 (19.9) | 16.3 (20.9) | 16.6 (24.9) |
| CC ½ (%)[1] | 91.9 (83.8) | 93.1 (89.2) | 94.4 (89.5) | 94.8 (90.4) | 94.8 (87.2) |
| $I/\sigma I$[1] | 9.47 (3.04) | 9.70 (2.88) | 9.14 (2.35) | 8.28 (1.87) | 7.34 (1.46) |
| Completeness (%)[1] | 100.0 (100.0) | 100.0 (100.0) | 100.0 (100.0) | 100.0 (100.0) | 100.0 (100.0) |
| Multiplicity[1] | 43.2 (29.7) | 51.2 (35.5) | 57.8 (39.9) | 58.8 (39.0) | 56.5 (35.0) |
| Unique Reflections[1] | 16,961 (855) | 16,959 (863) | 16,966 (866) | 16,966 (865) | 16,971 (861) |
| Wilson B-factor ($Å^2$) | 18.52 | 20.31 | 22.63 | 24.80 | 26.59 |

| **Data Collection** | **132 kGy** | **154 kGy** | **186 kGy** | **208 kGy** | **220 kGy** |
|---|---|---|---|---|---|
| Number of Integrated Lattices | 8,546 | 7,476 | 6,825 | 6,297 | 5,738 |
| Number of Merged Lattices | 8,512 | 7,442 | 6,796 | 6,255 | 5,699 |
| Unit Cell a, b, c (Å) | 39.4, 78.4, 48.1 | 39.4, 78.4, 48.1 | 39.4, 78.4, 48.1 | 39.4, 78.3, 48.1 | 39.4, 78.3, 48.1 |
| β (°) | 97.7 | 97.7 | 97.7 | 97.7 | 97.7 |
| Resolution, overall (Å) | 40.70 – 2.10 | 40.71 – 2.10 | 40.72 – 2.10 | 40.72 – 2.10 | 40.73 – 2.10 |
| Resolution, high (Å) | 2.14 – 2.10 | 2.14 – 2.10 | 2.14 – 2.10 | 2.14 – 2.10 | 2.14 – 2.10 |
| $R_{split}$[1] | 16.8 (28.7) | 17.7 (39.5) | 18.4 (51.2) | 18.5 (61.8) | 19.6 (76.9) |
| CC ½ (%)[1] | 95.3 (84.3) | 94.9 (75.0) | 94.5 (63.6) | 95.6 (58.4) | 95.6 (47.1) |
| $I/\sigma I$[1] | 6.28 (1.15) | 5.19 (0.92) | 4.23 (0.77) | 3.43 (0.70) | 2.88 (0.63) |
| Completeness (%)[1] | 100.0 (100.0) | 100.0 (100.0) | 100.0 (100.0) | 100.0 (100.0) | 100.0 (100.0) |
| Multiplicity[1] | 50.6 (28.0) | 44.9 (20.8) | 41.4 (15.5) | 38.1 (10.9) | 34.3 (8.2) |
| Unique Reflections[1] | 16,977 (862) | 16,981 (860) | 16,983 (856) | 16,986 (859) | 16,989 (861) |
| Wilson B-factor ($Å^2$) | 27.84 | 29.60 | 30.88 | 31.55 | 37.88 |

[1]High resolution statistics in parentheses

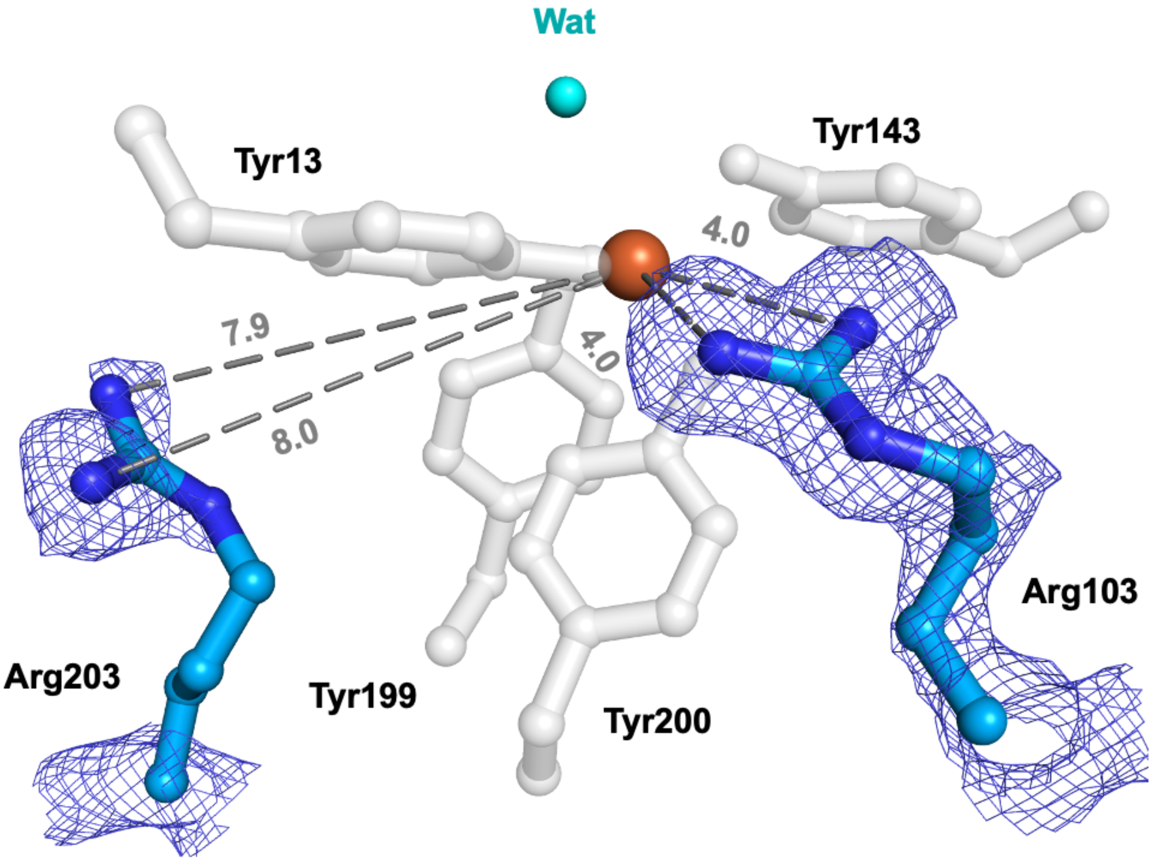

**Figure S1.** The iron center of FutA (Fe(III) state) by neutron diffraction. The nuclear density ($2F_{obs} - F_{calc}$, blue, contoured at 1.5σ) reveals positioning of the side chain of Arg103 close to the tyrosinates, while the Arg203 side chain does not engage in interactions (similar to the SFX structure **Fig. 2D**). Carbons shown blue, heteroatoms colored as in **Fig. 1**.

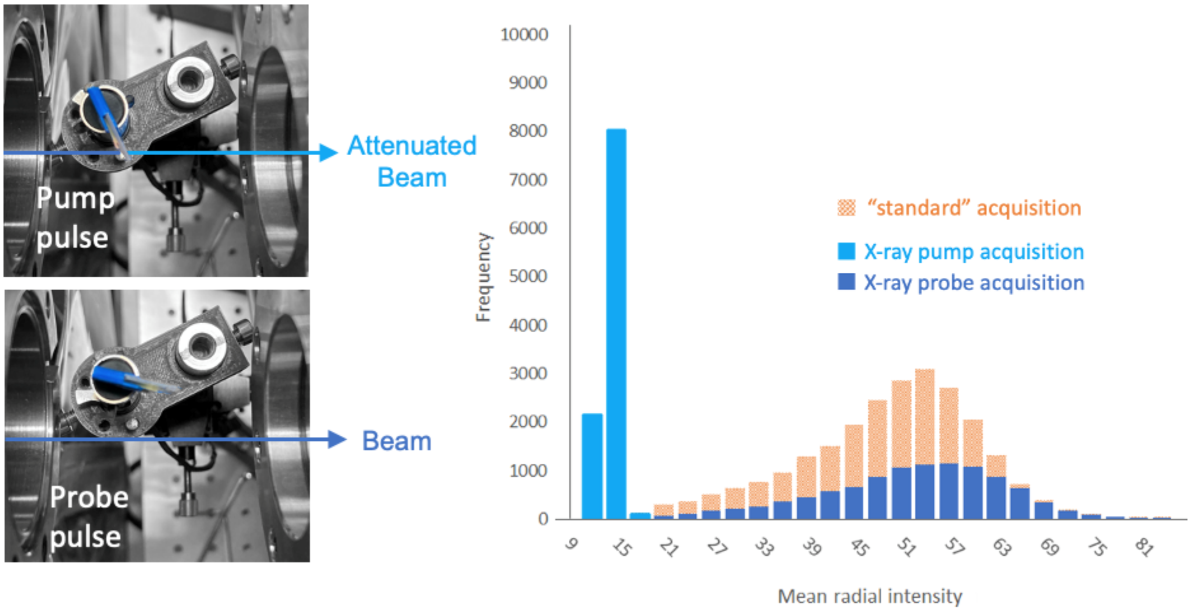

**Figure S2:** Overview of the experimental setup and data analysis for the XFEL X-ray pump probe experiment. A sapphire flipper-attenuator was placed in the path of the XFEL beam that was TTL triggered from a signal generator to move the wafer with alternating pulses, while two diffraction images corresponding to X-ray pump and X-ray probe were collected (left). The average diffraction intensity (diffraction spots and background) was calculated for each diffraction image and plotted as a histogram against frequency (right). The X-ray pump (light blue) is distinguished from the X-ray probe (dark blue) by its lower average diffraction intensity. For comparison, a histogram for a standard SFX experiment is shown (salmon).

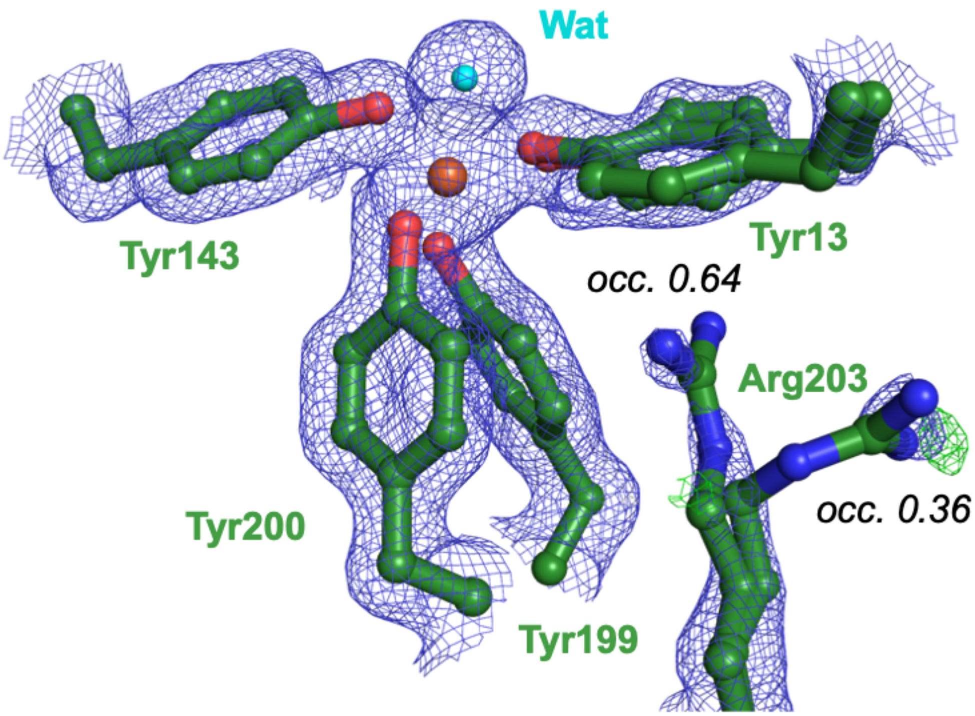

**Figure S3.** Refined SFX X-ray probe structure **Fig. 4**. The side chain of Arg203 was refined in dual occupancy, as indicated. Density shown in blue is 2Fo-Fc at 1.5 $\sigma$, difference density Fo-Fc, shown in green, 3 $\sigma$; no negative difference density was observed. Heteroatoms colored as in **Fig. 1**.

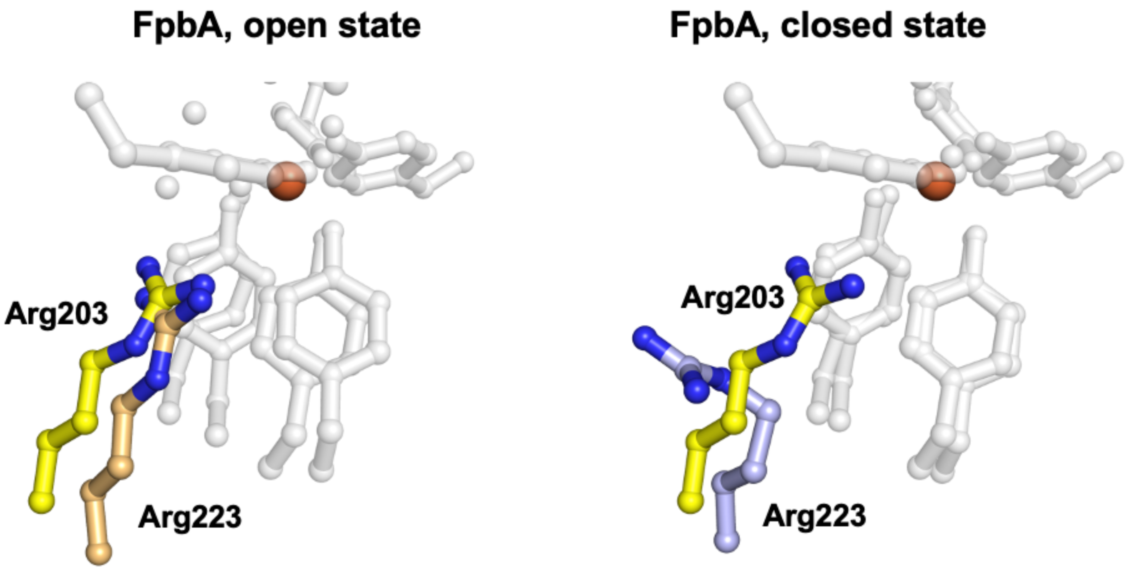

**Figure S4:** Overlay of Fe(II) bound *Prochlorococcus* MED4 FutA (yellow) with *Thermus thermophilus* FbpA in the open (light orange, PDB:3WAE) and closed confirmations (light purple, PDB:4ELR). As FpbA switches from an open to a closed conformation, a carbonate ion is lost and Arg233 is repositioned to maintain a net neutral charge in the binding site. The repositioning of Arg233 in FbpA is highly similar to the structural switch of Arg203 in FutA.

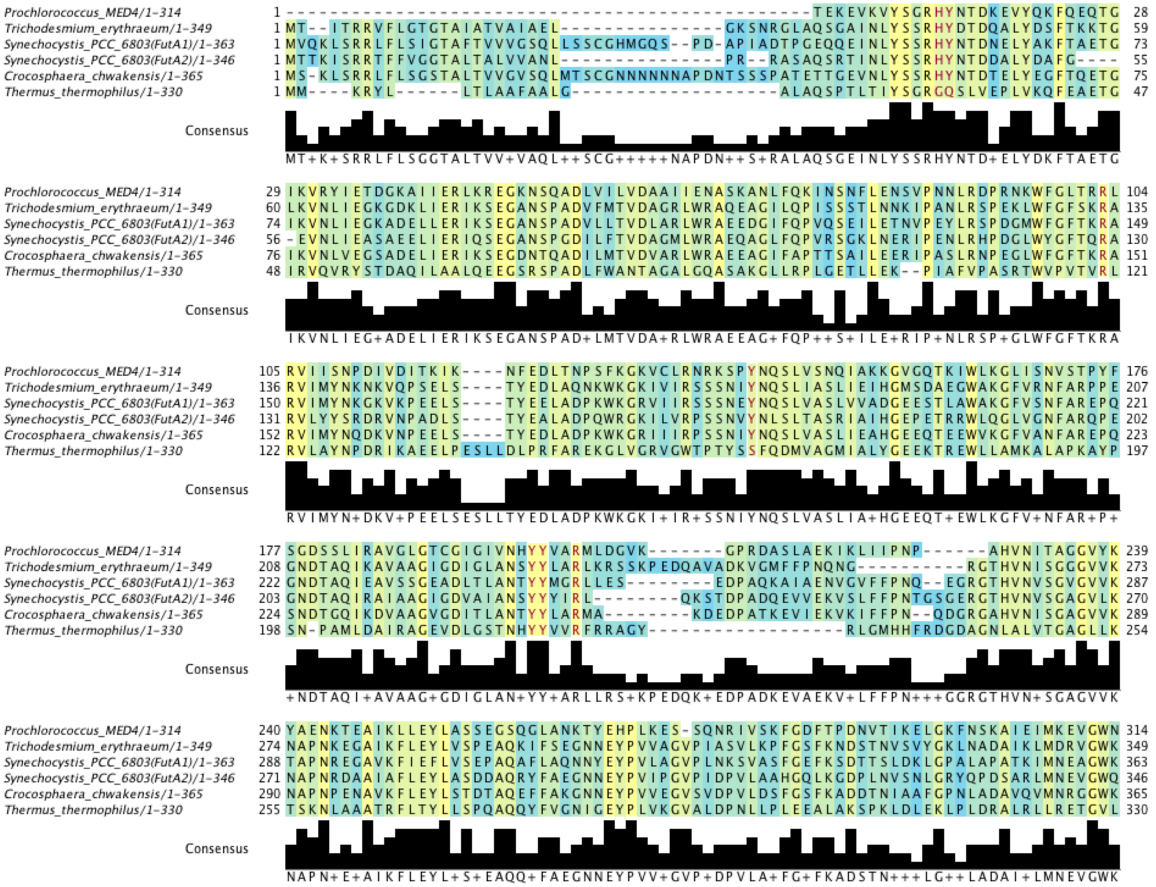

**Figure S5:** Multiple sequence alignment of the FutA homologues from *Prochlorococcus* MED4 (UniProt ID: Q7V0T9, excluding signal peptide amino acids 1-27), *Trichodesmium erythraeum* (UniProt ID: Q10Z45), *Synechocystis* PCC 6803 (FutA1 UniProt ID: P72827 and FutA2 UniProt ID: Q55835), *Crocosphaera chwakensis* (A3IPT8), and *Thermus thermophilus* (UniProt ID: Q5SHV2) carried out with Clustal Omega (26) and Jalview (27). Iron coordinating amino acids in Prochlorococcus MED4 are shown in red text. The consensus graph indicates the modal residue and the number of times the modal residue appears at each position. "+" denotes positions where the modal residue is shared by more than one amino acid type. The blue and yellow gradient shading indicates the degree of conservation of the physio-chemical properties of the amino acids at each position, where blue indicates low conservation and yellow indicates high conservation.

### References

1. J. J. Almagro Armenteros *et al.*, SignalP 5.0 improves signal peptide predictions using deep neural networks. *Nature Biotechnology* **37**, 420-423 (2019).
2. J. H. Beale *et al.*, Successful sample preparation for serial crystallography experiments. *Journal of Applied Crystallography* **52**, 1385-1396 (2019).
3. S. Horrell *et al.*, Fixed Target Serial Data Collection at Diamond Light Source. *J Vis Exp* 10.3791/62200 (2021).
4. W. Kabsch, XDS. *Acta Crystallographica Section D* **66**, 125-132 (2010).

5. P. R. Evans, G. N. Murshudov, How good are my data and what is the resolution? *Acta Crystallographica Section D* **69**, 1204-1214 (2013).
6. Z. Otwinowski, W. Minor, Processing of X-ray diffraction data collected in oscillation mode. *Methods in Enzymology* **276**, 307-326 (1997).
7. G. Winter *et al.*, DIALS: implementation and evaluation of a new integration package. *Acta Crystallographica Section D* **74**, 85-97 (2018).
8. M. Uervirojnangkoorn *et al.*, Enabling X-ray free electron laser crystallography for challenging biological systems from a limited number of crystals. *Elife* **4** (2015).
9. P. Howell, G. Smith, Identification of heavy-atom derivatives by normal probability methods. *Journal of applied crystallography* **25**, 81-86 (1992).
10. T. Nakane *et al.*, Data processing pipeline for serial femtosecond crystallography at SACLA. *Journal of Applied Crystallography* **49**, 1035-1041 (2016).
11. J. Hattne *et al.*, Accurate macromolecular structures using minimal measurements from X-ray free-electron lasers. *Nature Methods* **11**, 545-548 (2014).
12. A. Vagin, A. Teplyakov, MOLREP: an Automated Program for Molecular Replacement. *Journal of Applied Crystallography* **30**, 1022-1025 (1997).
13. P. Emsley, B. Lohkamp, W. G. Scott, K. Cowtan, Features and development of Coot. *Acta Crystallogr D Biol Crystallogr* **66**, 486-501 (2010).
14. G. N. Murshudov *et al.*, REFMAC5 for the refinement of macromolecular crystal structures. *Acta Crystallographica Section D* **67**, 355-367 (2011).
15. J. Sanchez-Weatherby *et al.*, Improving diffraction by humidity control: a novel device compatible with X-ray beamlines. *Acta Crystallogr D Biol Crystallogr* **65**, 1237-1246 (2009).
16. C. R. Harris *et al.*, Array programming with NumPy. *Nature* **585**, 357-362 (2020).
17. W. McKinney, Data Structures for Statistical Computing in Python. *Proceedings of the 9th Python in Science Conference* **445**, 56-61 (2010).
18. P. Virtanen *et al.*, SciPy 1.0: fundamental algorithms for scientific computing in Python. *Nature Methods* **17**, 261-272 (2020).
19. J. D. Hunter, Matplotlib: A 2D Graphics Environment. *Computing in Science & Engineering* **9**, 90-95 (2007).
20. G. N. Murshudov *et al.*, REFMAC5 for the refinement of macromolecular crystal structures. *Acta Crystallogr D Biol Crystallogr* **67**, 355-367 (2011).
21. L. Catapano *et al.*, Neutron crystallographic refinement with REFMAC5 of the CCP4 suite. *bioRxiv* 10.1101/2023.08.13.552925, 2023.2008.2013.552925 (2023).
22. S. Grazulis *et al.*, Crystallography Open Database (COD): an open-access collection of crystal structures and platform for world-wide collaboration. *Nucleic Acids Res.* **40**, D420-427 (2012).
23. C. S. Bury, J. C. Brooks-Bartlett, S. P. Walsh, E. F. Garman, Estimate your dose: RADDOSE-3D. *Protein Science* **27**, 217-228 (2018).
24. J. L. Dickerson, P. T. N. McCubbin, E. F. Garman, RADDOSE-XFEL: femtosecond time-resolved dose estimates for macromolecular X-ray free-electron laser experiments. *J. Appl. Cryst.* **53**, 549-560 (2020).
25. J. L. Dickerson, E. F. Garman, Doses for experiments with microbeams and microcrystals: Monte Carlo simulations in RADDOSE-3D. *Protein Sci* **30**, 8-19 (2021).
26. F. Sievers *et al.*, Fast, scalable generation of high-quality protein multiple sequence alignments using Clustal Omega. *Molecular Systems Biology* **7**, 539 (2011).
27. A. M. Waterhouse, J. B. Procter, D. M. A. Martin, M. Clamp, G. J. Barton, Jalview Version 2—a multiple sequence alignment editor and analysis workbench. *Bioinformatics* **25**, 1189-1191 (2009).
28. D. Polyviou *et al.*, Structural and functional characterization of IdiA/FutA (Tery_3377), an iron-binding protein from the ocean diazotroph Trichodesmium erythraeum. *Journal of Biological Chemistry* **293**, 18099-18109 (2018).
29. P. Lu *et al.*, Functional characterisation of two ferric-ion coordination modes of TtFbpA, the periplasmic subunit of an ABC-type iron transporter from Thermus thermophilus HB8. *Metallomics* **11**, 2078-2088 (2019).
30. S. Wang *et al.*, A novel mode of ferric ion coordination by the periplasmic ferric ion-binding subunit FbpA of an ABC-type iron transporter from Thermus thermophilus HB8. *Acta Crystallographica Section D* **70**, 196-202 (2014).
31. C. J. Williams *et al.*, MolProbity: More and better reference data for improved all-atom structure validation. *Protein Sci* **27**, 293-315 (2018).
